# Quanta microscopy enables fluorescence imaging at the native bit-plane rate

**DOI:** 10.64898/2026.10.26.761986

**Authors:** Jooyeong Yun, Keyi Han, Aleksandra Janusz-Kaminska, Wenhao Zhang, Zhi Ling, Yan Huang, Chunhui Xu, Gary J. Bassell, Shu Jia

## Abstract

Live-cell fluorescence microscopy remains constrained by limited photon budgets and exposure-integrated detection, imposing a fundamental trade-off between temporal sampling and image quality. Here we present quanta microscopy (QM), an optical–computational framework that integrates high-speed single-photon acquisition with self-supervised spatiotemporal reconstruction. QM transforms highly sparse binary photon-detection maps, or bit planes, into multibit grayscale fluorescence image sequences, generating one output frame per bit plane at the native 50 kHz sensor cadence without temporal binning. We demonstrate QM across biological specimens spanning diverse spatial and temporal characteristics, including subcellular organelles with distinct morphologies, rapid contractile dynamics in beating hiPSC-derived cardiomyocytes, and electrically stimulated calcium-response transients in primary cortical neurons. By uniting quanta-level acquisition with data-driven image formation, QM broadens the scope of live-cell fluorescence microscopy for quantitative analysis of rapid cellular physiology under photon-limited conditions.

## INTRODUCTION

Fluorescence microscopy enables molecularly specific visualization of cellular organization and function in living systems^1–3^. Its performance, however, is fundamentally constrained by the finite photon budget of fluorescent probes. Photon collection can be enhanced by increasing the excitation intensity or extending the exposure time; however, the former exacerbates photobleaching and phototoxicity, whereas the latter introduces temporal averaging and motion blur^4,5^. Consequently, imaging rapid cellular dynamics requires short acquisition windows that preserve temporal information while yielding photon-sparse measurements.

To date, instrumental and computational advances have sought to mitigate this limitation. High-speed camera architectures, including optical multiplexing and pixel-programmable acquisition, can accelerate readout or balance signal-to-noise ratio against frame rate across the field of view^6,7^. Meanwhile, deep-learning methods can denoise photon-starved fluorescence images^8–10^ or interpolate intermediate states to increase the apparent temporal sampling rate^11^. Most of these approaches, however, operate on conventional exposure-integrated measurements, in which photocharge is accumulated over a fixed interval before digitization, collapsing the temporal structure of photon arrivals within each pixel into a single intensity value. Subsequent computation can enhance or infer image content but may not directly recover temporal information that was not retained during acquisition.

Quanta image sensors offer an alternative acquisition regime by recording single-photon events at high frame rates as binary maps or photon counts^12–15^. Single-photon avalanche diode (SPAD) arrays, for example, generate sparse 1-bit photon-detection maps, or bit planes, at a cadence determined by the sensor readout cycle^16–18^. Furthermore, computational methods have enabled the reconstruction of intensity images and dynamic scenes from such photon-limited measurements^19–33^. However, quanta imaging remains underexplored in live-cell fluorescence microscopy, in which low and spatially heterogeneous photon flux must be reconciled with the need to preserve fine subcellular structures and rapid intensity transients. It therefore remains unclear whether grayscale image sequences can be reconstructed at the native bit-plane rate without temporal binning while preserving biologically relevant spatial and temporal information.

Here, we introduce quanta microscopy (QM), an integrated fluorescence imaging framework that combines high-speed single-photon acquisition with self-supervised spatiotemporal reconstruction. QM produces one grayscale image for every acquired bit plane without temporal binning, thereby preserving the native sampling cadence while leveraging learned spatial and temporal correlations to derive fluorescence intensity from sparse photon-detection events. We demonstrate QM across biological specimens spanning diverse spatial and temporal characteristics, including subcellular organelles with distinct morphologies, rapid contractile dynamics in live cardiomyocytes, and electrically evoked calcium transients in individual neurons. Together, these results establish QM as a practical and versatile platform for high-speed fluorescence microscopy under photon-limited conditions.

## RESULTS

### Principle and framework of QM

In conventional fluorescence microscopy, charge-integrating cameras accumulate photoelectrons over a fixed exposure and digitize the resulting charge as multibit pixel values representing the time-integrated fluorescence signal. Under photon-limited conditions, increasing the image signal-to-noise ratio generally requires longer exposures or temporal averaging, both of which blur the signal in time and can obscure rapid biological dynamics (**Fig. 1a**).

**Figure 1.**
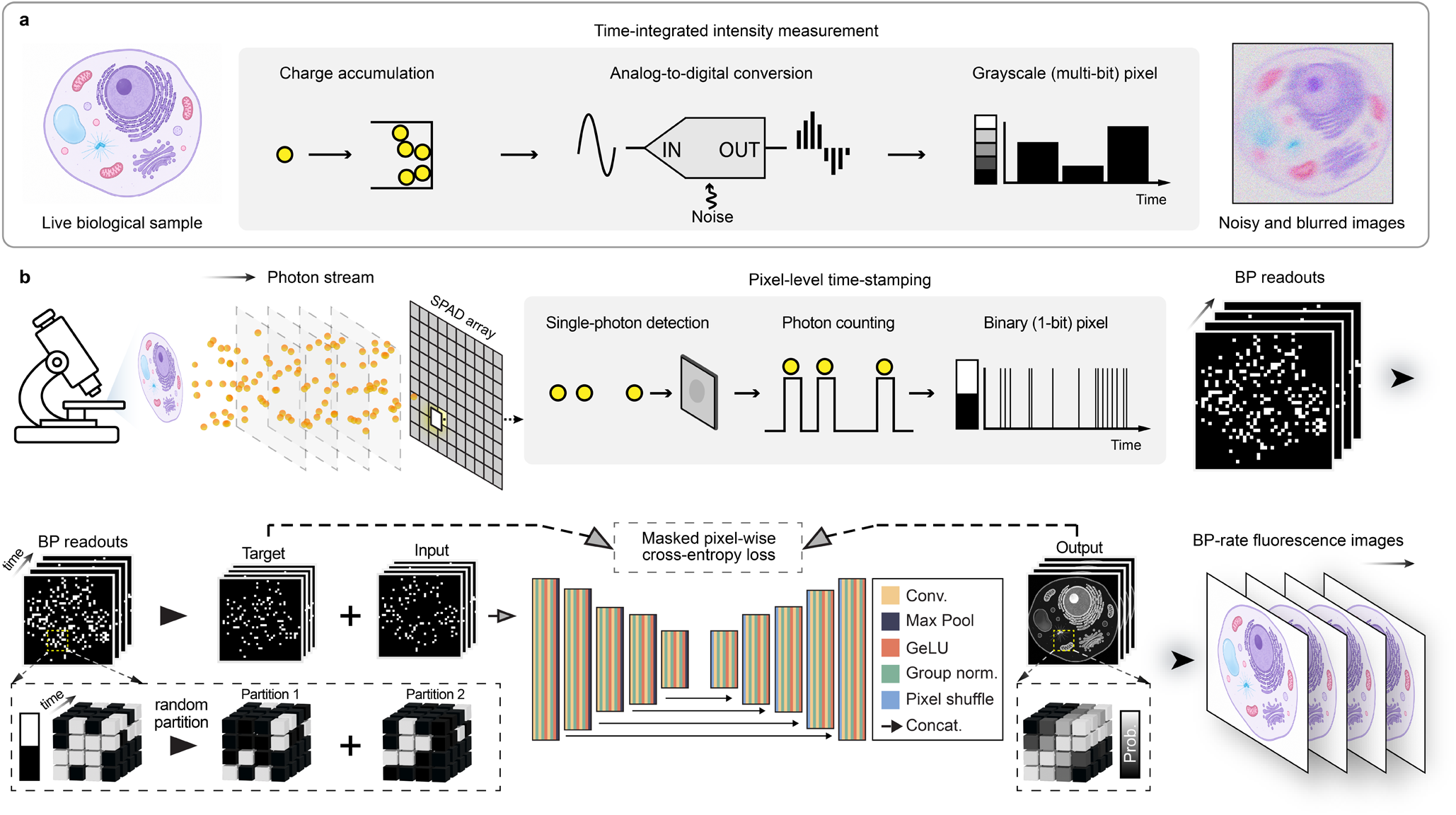
Principle and framework of quanta microscopy (QM). **(a)** Conventional exposure-integrated fluorescence imaging. Detected fluorescence photons are converted to photoelectrons, accumulated during a fixed exposure, and digitized as multibit grayscale pixel values. Under photon-limited or high-speed conditions, insufficient photon collection and readout noise reduce signal-to-noise ratio, whereas temporal integration can blur rapid dynamics. **(b)** Workflow of QM. An SPAD array records fluorescence from biological specimens as a temporally ordered sequence of sparse 1-bit photon-detection maps, termed bit planes (BPs). During self-supervised training, detection events within spatiotemporal BP volumes are randomly partitioned into complementary input and target volumes, and a hybrid 3D/2D ResUNet is optimized using a masked voxel-wise cross-entropy loss. During inference, the network transforms successive BP volumes into a frame-aligned multibit grayscale fluorescence image sequence, yielding one output frame per BP at the native BP cadence without temporal binning.

QM instead combines high-speed single-photon acquisition with self-supervised spatiotemporal reconstruction to generate one grayscale fluorescence image for each acquired bit plane, without temporal binning (**Fig. 1b**). The platform comprises coupled acquisition and computational modules (**Methods** and **Supplementary Fig. S1**). We implemented the acquisition module on a conventional epi-fluorescence microscope equipped with a 100×, 1.45NA objective and a SPAD camera operating at a frame rate of 50 kHz (**Fig. 1b, top** and **Methods**). In Geiger mode, each SPAD pixel reports whether at least one photon-detection event occurred during a readout interval^18^. Each camera readout therefore forms a bit plane (BP), a temporally indexed 1-bit map of pixels registering detection events. Although the BP sequence retains the native acquisition cadence of the SPAD camera, each BP is highly sparse in nature and does not directly represent a conventional grayscale fluorescence image. The reconstruction module converts the BP sequence into a frame-aligned grayscale sequence, using neighboring BPs as spatiotemporal context rather than summing them.

For reconstruction, we adapted a self-supervised photon-splitting strategy^20^ to spatiotemporal BP data, using a hybrid 3D/2D ResUNet (**Methods** and **Supplementary Note 1**). During training, BP stacks were randomly cropped into space–time volumes containing 32 or 64 consecutive BPs, corresponding to temporal windows of 0.64 or 1.28 ms at the 50 kHz acquisition rate. Detection events within each crop were partitioned into complementary input and target volumes, with the splitting probability treated as a training hyperparameter (**Fig. 1b, bottom**). The network was optimized using a masked voxel-wise cross-entropy loss that accounted for the dependence between the paired binary partitions (**Methods**). Temporal crop depth, photon-splitting probability, and BP training-pool size were adjusted for datasets with different photon levels and temporal dynamics (**Supplementary Tables S1** and **S2**). During inference, the network predicted frame-aligned fluorescence intensity from successive BP volumes while preserving the input frame count and temporal indexing. QM therefore produced one grayscale frame per BP at the native 50 kHz cadence, corresponding to a 20 µs sampling interval without temporal binning. This acquisition and reconstruction cadence was maintained throughout the following biological demonstrations.

### QM reconstructs diverse subcellular structures in HeLa cells

We evaluated QM on subcellular targets spanning diverse morphologies and signal levels. To quantify BP measurement sparsity, we calculated mean detection density as the average number of registered detection events per pixel per BP (**Methods** and **Supplementary Fig. S2**). We first imaged SYTO-16-labeled nuclei in fixed HeLa cells. At a mean detection density of 0.0034 detections per pixel per BP, nuclear morphology remained indiscernible in individual BPs (**Fig. 2a**). Following self-supervised training on 5,000 consecutive BPs collected over 100 ms, QM recovered nuclear boundaries and intranuclear intensity variations, consistent with those observed in the sCMOS reference images acquired with a 100-ms exposure (**Fig. 2b-d**). We next examined immunostained microtubules, whose fine filamentous architecture posed a more stringent reconstruction challenge. Here, microtubules exhibited significantly lower mean detection density (0.0029), corresponding to only 760 detection events per BP, making their fine filamentous structures particularly challenging to reconstruct (**Fig. 2e**). Nevertheless, QM resolved continuous microtubule filaments with clear contrast, whereas the corresponding sCMOS image remained strongly noise-limited (**Fig. 2f-j**).

**Figure 2.**
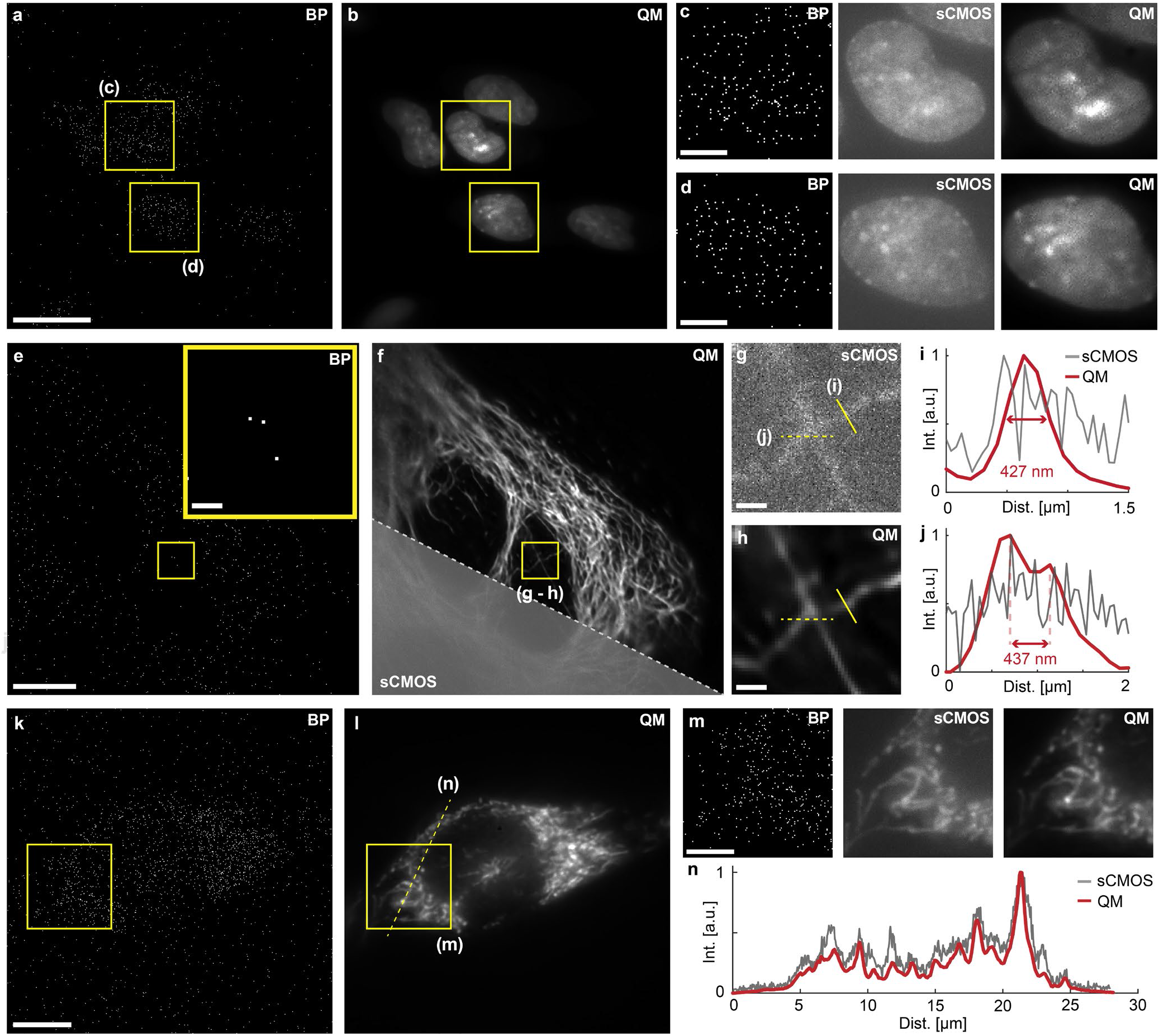
Imaging subcellular structures in fixed and live HeLa cells. **(a, b)** Representative BP (a) and corresponding QM reconstruction (b) of SYTO-16-labeled nuclei in fixed HeLa cells. **(c, d)** Zoomed-in BP, sCMOS, and QM images of the boxed regions in (a, b). **(e)** Representative BP of immunostained microtubules in fixed HeLa cells. Inset, zoomed-in view of the boxed region. **(f)** Diagonal split-view of the QM (upper right) and corresponding sCMOS reference (lower left) images. **(g, h)** Zoomed-in sCMOS (g) and QM (h) images of the boxed region in (f). **(i, j)** Cross-sectional intensity profiles along the solid (i) and dashed (j) lines as indicated in (g, h). Red double-headed arrows in (i, j) measure the corresponding QM intensity profiles. **(k, l)** Representative BP (k) and corresponding QM (l) images of MitoTracker Green-labeled mitochondria in live HeLa cells. **(m)** Zoomed-in BP, sCMOS reference, and QM images of the boxed regions in (k, l). **(n)** Intensity profiles along the dashed line in (l), exhibiting high-contrast subcellular structures revealed by QM. All BP data were acquired at 50 kHz, and all sCMOS reference images were acquired with 100 ms exposures. Scale bars: 20 µm (a), 10 µm (e, k), 5 µm (c, d, m), 1 µm (inset in e, g, h).

Moreover, we imaged MitoTracker-labeled mitochondria in live HeLa cells with a mean detection density of 0.0090. QM recovered the heterogeneous mitochondrial network and revealed its local intensity variations (**Fig. 2k, l**). Compared with the corresponding sCMOS images, the QM reconstructions showed reduced background fluctuations while preserving mitochondria-associated intensity features (**Fig. 2m, n**). Despite operating at mean detection densities below 0.01 detections per pixel per BP, an exceptionally photon-sparse regime compared with representative low-light natural-scene and fluorescence-imaging studies^9,20,32^, QM underwent the self-supervised reconstruction of morphologically diverse subcellular structures from both fixed- and live-cell fluorescent probes, including under conditions in which conventional sCMOS imaging remained noise-limited.

### QM resolves rapid contractile dynamics in hiPSC-derived cardiomyocytes

Human induced pluripotent stem cell-derived cardiomyocytes (hiPSC-CMs) provide a physiologically relevant model for cardiac disease modeling and pharmacological testing^34–36^. Cardiomyocyte contraction produces coordinated deformation across the entire cell along with rapid subcellular motion, requiring both wide-field capture and high temporal sampling^37,38^. Under photon-limited fluorescence conditions, these multiscale dynamics are often compromised for simultaneous observation.

Here, we evaluated QM in beating hiPSC-CMs labeled with tetramethylrhodamine methyl ester, a mitochondrial membrane-potential dye (**Methods**). Individual BPs did not reveal the mitochondrial network, whereas QM recovered heterogeneous mitochondrial structures and fine subcellular features (**Fig. 3a** and **Supplementary Fig. S3**). By generating one reconstructed frame per BP, QM sampled the contraction–relaxation cycles at 20 µs intervals, corresponding to a 50 kHz output cadence and thereby a two-to-three orders of magnitude reduction in frame interval relative to conventional epifluorescence imaging (**Fig. 3b**). Associated morphological dynamics were analyzed using frame-wise decorrelation relative to the initial frame (**Methods** and **Supplementary Note 2.1**). Notably, the resulting trace clearly delineated the contraction and relaxation phases (**Fig. 3c** and **Supplementary Fig. S4**), while overlays of the first and last frames within each phase revealed the corresponding subcellular spatiotemporal displacements (**Fig. 3d**). In contrast, such spatiotemporal dynamics are not readily resolvable with conventional imaging without sacrificing either sensor readout area or signal-to-noise ratio^6,7,39^.

**Figure 3.**
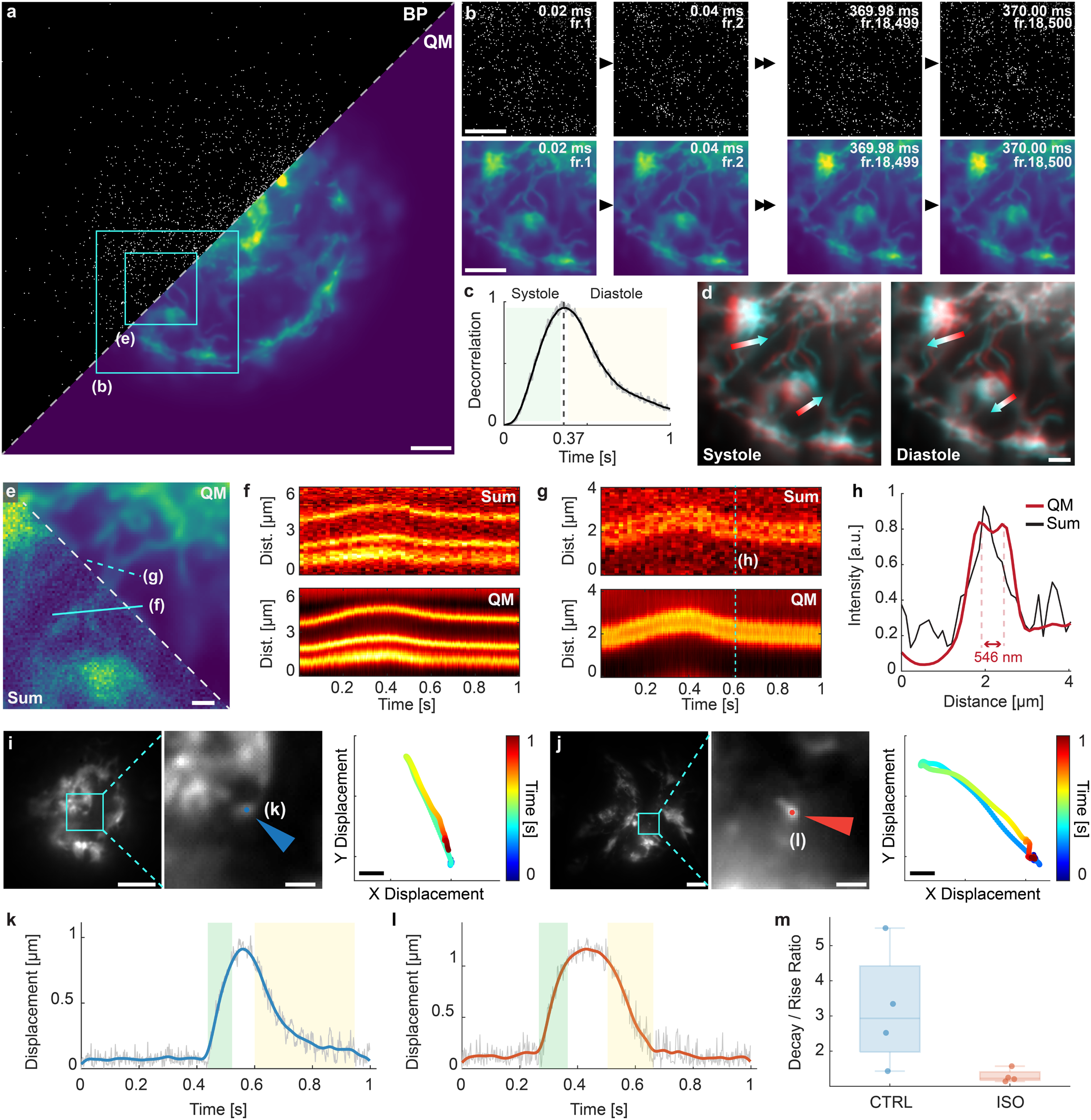
Imaging contractile dynamics in hiPSC-derived cardiomyocytes. **(a)** Diagonal split-view comparison of a representative BP and corresponding QM images of TMRM-labeled beating cardiomyocytes. **(b)** Zoomed-in representative BPs (top) and corresponding QM images (bottom) of the boxed region in (a) at the indicated time points during a contraction–relaxation cycle. **(c)** Frame-wise decorrelation relative to the initial QM frame over a single cycle, separating phases of systole and diastole at the maximum decorrelation at *t* = 0.37 s. Gray and black traces denote the raw and smoothed decorrelation values, respectively. **(d)** Overlays of the first and last QM frames within the systolic and diastolic phases, revealing representative subcellular displacements (arrows). **(e)** Diagonal split-view comparison of the QM (upper right) and temporally summed (lower left) images from the boxed region in (a). The Sum image was generated by integrating 1,000 consecutive BPs over 20 ms. **(f, g)** Kymographs of Sum (top) and QM (bottom) along the solid (f) and dashed (g) lines in (e). **(h)** Intensity profiles at the time point marked by the vertical dashed line in (g), in which QM resolved a 546-nm separation between adjacent mitochondrial structures. **(i, j)** Representative QM image (left), zoomed-in view of the boxed region (middle), and corresponding time-coded x–y displacement trajectories (right) for untreated control (i) and isoproterenol-treated (ISO; j) cardiomyocytes (arrows). **(k, l)** Sub-micrometer displacement traces of the components indicated in (i, j). Gray and colored traces show the raw measurements and smoothed trajectories, respectively. Shaded regions indicate the rise (green) and decay (yellow) intervals used for kinetic analysis. **(m)** Boxplot of the decay-to-rise time ratio for control (CTRL) and ISO-treated cardiomyocytes (*n* = 4 per group). All BPs were acquired at 50 kHz, with one QM frame reconstructed per BP. Scale bars: 5 µm (a, b), 2 µm (d, middle panels of i, j), 1 µm (e), 10 µm (left panels of i, j), 0.2 µm (trajectory plots in i, j).

Next, we compared QM-reconstructed sequences with images generated by temporally summing consecutive BPs (**Fig. 3e**). Summing 1,000 BPs over a 20 ms integration window pooled detection events and improved signal visibility, which, however, blurred rapid motion and attenuated transient structural changes (**Fig. 3f, g**). By contrast, QM reduced shot-noise fluctuations while preserving the native BP cadence, thereby resolving continuous mitochondrial motion and fine subcellular details without compromising signal quality (**Fig. 3h** and **Supplementary Video 1**).

Furthermore, the use of QM allows us to precisely differentiate the variations in contractility between the control group and the group administered isoproterenol, a compound known to increase heart rate and contractility and used therapeutically for bradycardia and other cardiac conditions^40–42^ (**Methods**). Subcellular landmarks were tracked throughout individual contraction cycles to obtain displacement trajectories (**Fig. 3i, j, Supplementary Fig. S5** and **Supplementary Note 2.2**). Untreated cardiomyocytes exhibited asymmetric trajectories, characterized by longer decay than rise phases and greater variability in the decay-to-rise time ratio (**Fig. 3k**). Relative to untreated cardiomyocytes, isoproterenol-treated cells typically showed more symmetric trajectories, with comparable rise and decay times and reduced variability in the decay-to-rise time ratio (**Fig. 3l, m**, and **Supplementary Videos 2, 3**). These differences are consistent with previously reported changes in contraction–relaxation kinetics induced by isoproterenol treatment^40,43^. Together, these results demonstrate QM as a platform for high-speed quantitative analysis of cardiomyocyte function, with potential applications in cardiac disease modeling and pharmacological testing.

### QM maps electrically evoked calcium dynamics in primary cortical neurons

Neuronal calcium signals are rapid and spatially heterogeneous, particularly within fine cellular processes, requiring fluorescence imaging with both high sensitivity, spatial resolution, and temporal sampling^44–47^. We therefore evaluated QM in primary cortical neurons loaded with Fluo-4 and locally stimulated using a custom electrical probe (**Fig. 4a, Methods** and **Supplementary Fig. S6**). Electrical stimulation elicited transient increases in neuronal fluorescence (**Fig. 4b, c** and **Supplementary Video 4**).

**Figure 4.**
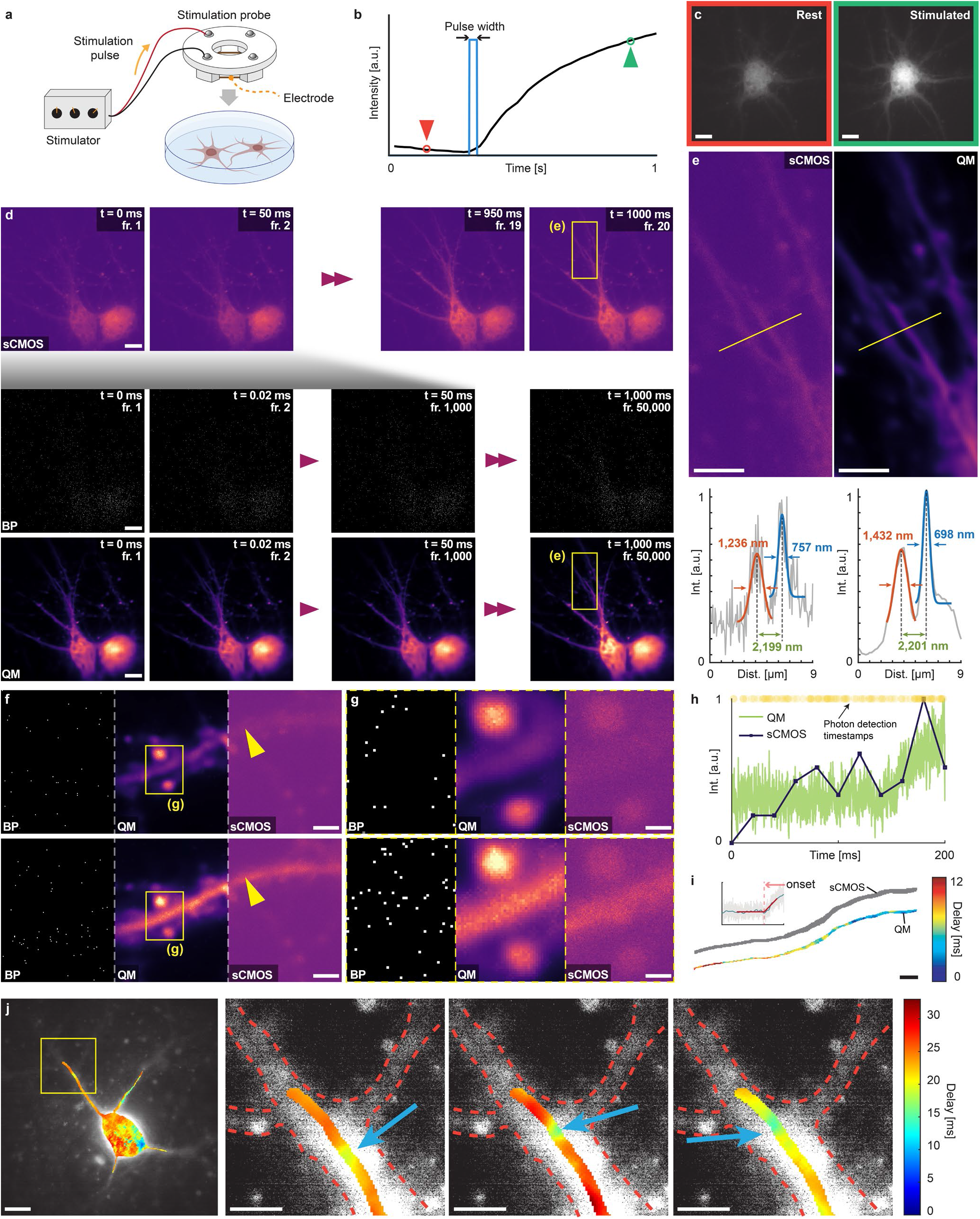
Mapping electrically evoked calcium dynamics in primary cortical neurons. **(a)** Schematic of the electrical stimulation setup for Fluo-4-loaded neurons. **(b)** Schematic of the stimulation pulse and associated neuronal fluorescence response. Blue lines mark the pulse width; red and green arrowheads indicate the resting and stimulated states shown in (c), respectively. **(c)** Representative fluorescence images of a neuron at rest (left) and following a 3 ms stimulation pulse (right). **(d)** Representative 1 s time-lapse sequences of neurons at DIV 8 from beam-split SPAD-sCMOS acquisition with a 300 ms stimulation pulse, showing sCMOS images (top; 20 Hz), raw BPs (middle; 50 kHz), and corresponding reconstructed QM images (bottom). **(e)** Zoomed-in sCMOS and QM images of the boxed region in (d). Bottom panel: corresponding intensity profiles along the yellow lines, indicating consistent measurements of feature widths and interfeature spacing. **(f)** Representative BP, QM, and sCMOS images before (top) and after (bottom) a 30 ms stimulation pulse, showing the calcium response along an individual axonal process at DIV 21. **(g)** Zoomed-in images of the boxed regions in (f). **(h)** Single-pixel fluorescence traces from the QM and sCMOS sequences, as indicated by the arrowheads in (f), together with the corresponding BP detection-event timestamps. **(i)** Calcium-response onset-delay maps along the axon derived from QM (color-coded by delay) and simultaneous 50 Hz sCMOS data (no resolvable delay variation). Inset, onset estimation from a spline-smoothed QM intensity trace using a rectified-linear fit. **(j)** QM-based onset-delay mapping of a complex neuronal network at DIV 16 across three 3 ms stimulation trials. Left, onset-delay map overlaid on the sCMOS image; right, zoomed-in images of the boxed branch region. Blue arrows indicate earlier branch-proximal response onsets, and red dashed lines outline adjacent neurites. All BPs were acquired at 50 kHz, with one QM frame reconstructed per BP. Scale bars: 10 µm (c, d, j), 5 µm (e, f, i), 2 µm (g).

Leveraging beam-split SPAD-sCMOS acquisition, QM generated grayscale fluorescence frames at 50 kHz, 2,500-fold higher than the 20 Hz sCMOS cadence required for adequate signal under the same imaging conditions, while preserving comparable signal levels and spatial features (**Fig. 4d, Supplementary Video 5** and **Supplementary Fig. S1**). As a result, this substantially enhanced spatiotemporal sampling enabled detailed mapping of calcium-response propagation along individual axons across distinct neuronal activity states (**Fig. 4e-g, Supplementary Video 6** and **Supplementary Fig. S7**). Furthermore, at the single-neuronal-pixel level, the stimulus-evoked fluorescence increase was difficult to identify directly from the sparse BP detections and was temporally undersampled in the conventional sCMOS trace (**Fig. 4h** and **Supplementary Fig. S8**). In contrast, QM recovered a continuous intensity transient without explicit spatiotemporal binning, and this pixelwise precision allowed us to identify the corresponding response onset at each pixel using QM-derived intensity traces (**Methods**). The resulting onset-delay map revealed a clear temporal gradient along the axon, consistent with directional propagation of the calcium response (**Fig. 4i, Supplementary Note 3.1** and **Supplementary Video 7**). In contrast, the same analysis applied to the corresponding sCMOS data acquired at 50 Hz produced a nearly uniform map because the conventional 20 ms frame interval was insufficient to resolve these latency differences (**Fig. 4i**). Finally, we extended QM-based onset mapping of neurons in a medium density monolayer culture, with overlapping dendrites and axons originating in neighboring cells (**Supplementary Fig. S9** and **Supplementary Video 8**). Across repeated stimulation trials, regions proximal to neuritic branch points exhibited calcium-response onsets 4.58 ± 1.16 ms earlier than those in adjacent distal segments, revealing a reproducible branch-associated latency difference that was not resolved by corresponding 50 Hz sCMOS imaging (**Fig. 4j** and **Supplementary Note 3.2**). Together, these results demonstrate that QM enables high-speed mapping of subcellular calcium dynamics and propagation latencies in complex neuronal morphologies, supporting studies of neuronal signal propagation and compartmentalized activity^48–50^.

## DISCUSSION

Live-cell fluorescence imaging remains constrained by limited photon budgets and exposure-integrated detection, imposing a fundamental trade-off between temporal sampling and image quality^51–53^. Overcoming this constraint is essential for resolving rapid cellular physiology without increasing excitation dose or temporally averaging transient signals. In this study, we present QM, an optical–computational framework that integrates high-speed single-photon acquisition with self-supervised spatiotemporal reconstruction. QM produces one multibit grayscale fluorescence frame per BP from highly sparse photon-detection measurements, retaining the native sensor cadence without temporal binning. QM recovered diverse subcellular structures, quantified rapid contractile dynamics in hiPSC-derived cardiomyocytes, and mapped millisecond-scale calcium-response latencies in complex neuronal morphologies. Although demonstrated here with epifluorescence, the platform could extend across optical architectures, combining quanta imaging at the native BP rate with volumetric context^54,55^, molecular multiplexing^56,57^, high-throughput screening^58,59^, and subdiffractional spatial resolution^60–62^. The platform could be further expanded through co-design and integration with meta-optics^63–65^, sensor hardware^66,67^, advanced computation^68,69^, and quantum-optical methods^70,71^. Furthermore, extending QM beyond single-cell imaging to complex cellular and tissue models^72–75^ could facilitate quantitative studies of multicellular physiology, disease modeling, and pharmacological screening. By uniting quanta-level acquisition with data-driven image formation, we anticipate QM to transform fluorescence microscopy, enabling fast, photon-limited, and light-sensitive biomedical research.

## METHODS

### Image acquisition setup

Widefield imaging experiments were performed using an epi-fluorescence microscope (Eclipse Ti2-U, Nikon Instruments). In the microscope system, we implemented a 100× oil-immersion objective lens with a numerical aperture of 1.45 (CFI Plan Apochromat Lambda 100× Oil, Nikon Instruments) and a piezo nano-positioner (Nano-F100S, Mad City Labs) for precise positioning. During the experiments, samples were excited using multicolor laser lines (405 nm, Coherent; 488 nm, 561 nm, 647 nm, MPB Communications), while the fluorescence was collected through a quadband dichroic mirror (ZT405/488/561/647, Chroma) and a corresponding emission filter (ZET405/488/561/647 m, Chroma). A micro-positioning system (MS2000, Applied Scientific Instrumentation) was used for accurate sample positioning. For image acquisition, an sCMOS camera (ORCA-Flash 4.0 V3, Hamamatsu Photonics, pixel size *P*_cam_ = 6.5 µm) and a SPAD camera (SPAD512S, Pi Imaging, pixel size *P*_cam_ = 16.38 µm) were used. All BP acquisitions used the intensity imaging mode of the SPAD camera with a fixed framerate of 50 kfps. For simultaneous image acquisition with both cameras, a beam-splitting setup was implemented at the camera output port of the microscope (**Supplementary Fig. S1**). The built-in 1.5× intermediate magnification changer of the Nikon microscope was used for selected experiments.

### QM network architecture

We employed a hybrid 3D/2D ResUNet architecture to reconstruct continuous fluorescence sequences from sparse 1-bit SPAD data (**Supplementary Note 1**). To prevent the temporal receptive field from expanding too rapidly at deeper layers, we restricted 3D convolutions to the first two network levels. This prevents a fully 3D network from becoming erroneously sensitive to zero-padded boundaries that physically mimic native data sparsity. The remaining deeper levels used strictly 2D convolutional blocks, mitigating padding artifacts while improving memory efficiency. For all experiments, the network depth was set to 5, with an initial feature size of 32 and a depth scale factor of 2. Structurally, the network comprised residual down-convolution and up-convolution blocks linked by skip connections to preserve high-resolution spatial details. Each down-convolution block contained three convolutional layers (with a residual addition between the first two) followed by max-pooling for down-sampling. Up-convolution blocks employed pixel shuffle for up-sampling, followed by residual convolutional refinement. Group normalization (8 groups) and GeLU activation functions were applied after each convolutional layer throughout the network.

### QM network training

Prior to training, the raw BP measurements were cleaned using a hot-pixel correction algorithm previously reported^20^. To reconstruct the underlying continuous signal from binary photon detection events, we applied a self-supervised approach. The sparse space-time volume was randomly partitioned into an input volume, X*_inp_*, and a target volume, X*_tar_*, via Bernoulli sampling. Because the binary nature of the data dictates that a photon assigned to the input cannot exist in the target, the input and target volumes are strictly correlated. To prevent the network from merely learning this deterministic complementarity, the loss function *L* was formulated as a masked voxel-wise cross-entropy loss

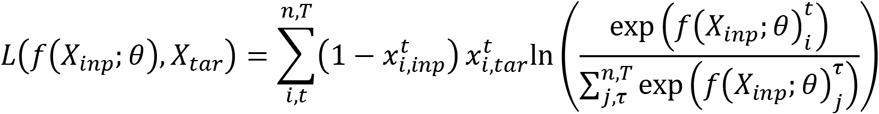

denotes the logit output of the 3D denoiser network parameterized by *θ*. The variables 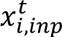 and 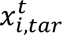 represent the binary pixel values at spatial index *i* and temporal index *t* within the input and target volumes, respectively. Similarly, 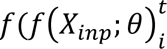 refers to the network’s logit prediction at the corresponding spatiotemporal coordinate. The critical masking term, 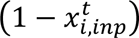, acts as the bitwise complement of the input image. By multiplying this term against the loss, the function explicitly excludes any pixels that contain a photon detection event in the input image. This effectively hides the complementary dependencies during backpropagation, allowing the model to accurately predict the underlying spatiotemporal photon arrival probability distribution. Models were trained using the ADAMW optimizer. To manage the rapid expansion of the receptive field in the 3D network and minimize the influence of zero-padding, large crop windows were maintained. Depending on the specific imaging conditions, training steps processed either four batches of 32 × 256 × 256 (TXY) random crops or two batches of 64 × 256 × 256 crops, with Bernoulli splitting probabilities (*p*) fixed or varied accordingly. While the total number of epochs varied across different cases, the overall number of training steps was kept similar to ensure consistent convergence. Computations were performed on NVIDIA RTX 3090 GPUs equipped with 24 GB of VRAM. Comprehensive details regarding case-specific crop sizes, batch configurations, epoch counts, *p*-value range, training data pool, and other tailored network hyperparameters are provided in **Supplementary Tables S1 and S2**. The overall training process generally required approximately 16–20 hours to reach convergence.

### QM network inference

Because the input recording is far longer than the temporal window the network processes at once, the full BP sequence is reconstructed patch-by-patch along the time axis. The volume is divided into consecutive spatiotemporal patches of size 64 × 512 × 512 (TXY), with adjacent patches overlapping by 48 frames. Within each overlap, the output was joined at the midpoint so that frames closer to the temporal center of each patch were retained, reducing boundary artifacts. The network output log-probabilities were exponentiated and normalized over each patch, then scaled by the total number of registered detection events in the corresponding input patch to restore the overall intensity level. The rescaled patches were subsequently stitched together to generate the full-length reconstructed sequence. All inference in this work used a patch size of 64 × 512 × 512 and a temporal overlap of 48 frames, except where a skip-inference scheme was used. For skip inference, the same 64 BP temporal window was centered on each sampled frame, and only the central reconstructed frame was retained. On a single NVIDIA RTX 3090 GPU, one 64 × 512 × 512 patch takes approximately 1.17 s in full precision

### Labeling mitochondria in live HeLa cells

Live mitochondria were stained and imaged in HeLa cells. Cells were cultured in DMEM with 10% FBS and 1% Penicillin-streptomycin (Pen-Strep) under 37 °C and 5% CO_2_ conditions. Once the cell cultures reached ∼80% confluency, cells were passaged and cultured in an imaging dish. On the day of imaging, cells were incubated in 50 nM MitoTracker Green (Thermo Fisher Scientific # M7514) in DMEM for 30 min. After removing the staining medium, the cells were washed 2 times with DMEM and immediately used for imaging.

### Labeling nucleus in fixed HeLa cells

Fixed nucleus staining was performed with HeLa cells. The culturing protocol was the same as described in the above section. A single coverslip was placed in a 35-mm culture dish containing gelatin and treated for 30 minutes. Subsequently, 2 mL DMEM was added, followed by the addition of 200 µL of HeLa cell suspension. The cells were allowed to grow and attach to the coverslip for 2–3 days. After incubation, the DMEM was removed from the culture dish. The labeling reagent was prepared with a concentration of 5 µM in a plastic tube by mixing 2 mL of FluoroBrite DMEM with 10 µL of SYTO 16 solution. The prepared labeling solution was prewarmed to 37 °C and then added to the culture dish. The cells were incubated for 1 hour at 37 °C under a CO₂ atmosphere. After labeling, the solution was removed, and the cells were washed three times with phosphate-buffered saline (PBS). For fixation, cells were treated with 4% paraformaldehyde (PFA) for 12 minutes inside the incubator. After fixation, the PFA was removed, and the cells were washed with PBS. The coverslip was carefully removed from the culture dish, and 25 µL of liquid antifade mountant was applied to a glass slide to minimize photobleaching. The coverslip was then placed onto the glass slide and secured with nail enamel, which was left to dry for 15 to 30 minutes before imaging.

### Immunostaining of microtubules in fixed HeLa cells

Fixed microtubule staining was performed with HeLa cells. The culturing protocol was the same as described in the above section. On the day of imaging, cells were first fixed with 0.3% (volume: volume) glutaraldehyde in extraction buffer and incubated for 1 minute at 37°C. The extraction buffer consists of 10 mM MES, 150 mM NaCl, 5 mM EDTA, 5 mM glucose, 5 mM MgCl2, and 0.25% (volume: volume) Triton X100 in ultrapure water. The buffer of the cells is then switched to 2% (volume: volume) glutar-aldehyde in cytoskeleton buffer at room temperature for 10 minutes. The cytoskeleton buffer is the afore-mentioned extraction buffer without the Triton X-100. Cells are then gently washed with blocking/per-meability (b/p) solution for 5 minutes, 3 times. The b/p solution consists of 2.5% (weight: volume) bovine serum albumin (BSA) and 0.1% (volume: volume) Triton X-100 in PBS. Then, the primary antibody that targets beta-tubulin was added to the cell dish with 2 mL of b/p solution at a concentration of 2 μg/mL (ThermoFisher, #32-2600). The cell dish was placed inside a humidified chamber at room temperature for 1 hour. After primary antibody tagging, the cell dish was washed with b/p solution 3 times, for 5 minutes each time. Then cells were co-labeled with 2 μg/mL of Goat anti-Mouse IgG conjugated with Alexa Fluor Plus 488 (ThermoFisher, #A32723) and 2 μg/mL of Goat anti-Mouse IgG conjugated with Alexa Fluor 647 (ThermoFisher, #A-21235). The staining took place inside a humidified chamber at room temperature for 1 hour. After secondary antibody labeling, the cell dish was washed with b/p solution 3 times for 5 minutes each and sequentially washed with PBS twice. The sample was finally stored in 2 mL of PBS solution for imaging.

### Acquisition, reconstruction, and signal-density analysis of HeLa-cell organelles

All sCMOS reference images were acquired with an exposure time of 100 ms. For each organelle dataset, QM was trained using 5,000 consecutive BPs acquired at 50 kHz. The selected BP sequence therefore spanned a total photon-collection interval of 100 ms, matching the nominal exposure duration of the corresponding sCMOS reference image and providing a duration-matched basis for comparison between the two imaging modalities.

BP sparsity was quantified using the mean detection density, defined as the number of photon detections per pixel per BP. Each BP contained 512×512 pixels, hence the mean detection density was calculated as

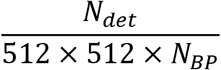

where *N_det_*is the total number of registered detection events and *N_BP_* is the number of analyzed BPs. For the signal-level comparison shown in **Supplementary Fig. S2**, the calculation was performed using the first 1,000 BPs from each acquisition.

### Preparation, staining, and imaging of hiPSC-CMs

The hiPSC line iPS (IMR90) (WiCell Research Institute) was maintained on Matrigel-coated 12-well plates in mTeSR1 medium (STEMCELL Technologies) with daily media changes. Cells were induced to differentiate into cardiomyocytes when they reached 85% confluence. Cells were treated with RPMI 1640 medium supplemented with 2% B27 minus insulin supplement (B27(-), ThermoFisher) (RPMI+B27(-)) containing 6 µM CHIR99021 (SelleckChem) for 48 h, followed by 24 h of RPMI+B27(-) to initiate cardiomyocyte differentiation. Subsequently, 5 µM IWR-1 (Sigma-Aldrich) was applied in RPMI+B27(-) for 48 h. Cultures were then maintained in RPMI supplemented with B27 plus insulin (B27(+), ThermoFisher) (RPMI+B27(+)), with media refreshed every 2-3 days until cardiomyocyte differentiation was evident.

On the day of imaging, mitochondria were labeled with tetramethylrhodamine methyl ester (TMRM; T668, ThermoFisher), a mitochondrial membrane potential-sensitive dye. The staining solution was prepared by diluting 2 µL of 100 µM TMRM stock solution in 2 mL of prewarmed culture medium, yielding a final concentration of 100 nM. The maintenance medium was replaced with the staining solution, and the cells were incubated for 45 min at 37 °C. The staining solution was then replaced with 2 mL of prewarmed maintenance medium, and the cells were allowed to recover for at least 30 min.

Before imaging, the maintenance medium was replaced with imaging medium consisting of FluoroBrite DMEM (Thermo Fisher Scientific, A18967-01) supplemented with 1% GlutaMAX (Thermo Fisher Scientific, 35050-061), 1% non-essential amino acids (NEAA; Thermo Fisher Scientific, 11140-050), and 2 g/L AlbuMAX (Thermo Fisher Scientific, 11020-021). During imaging, the imaging dish was maintained in a stage-top incubator at 37 °C, 5% CO₂, and approximately 95% relative humidity.

For pharmacological treatment, isoproterenol was diluted from a 100 mM stock solution to a final concentration of 100 µM in the imaging medium. TMRM-labeled hiPSC-CMs were incubated in the isoproterenol-containing imaging medium for 10 min and then imaged under the same environmental and acquisition conditions as the untreated control group.

### Cardiomyocyte morphological dynamics analysis

Cardiomyocyte morphological dynamics were quantified from QM-reconstructed image sequences using frame-wise decorrelation and subcellular feature tracking. Each frame was compared with the initial frame using the two-dimensional Pearson correlation coefficient, and decorrelation was defined as *D*(*t*) = 1 − *ρ*(*t*). The normalized decorrelation trace was smoothed using a moving mean, and the time of maximum decorrelation was used to separate the systolic and diastolic phases.

For subcellular motion analysis, representative dot-like features were manually selected and tracked by fitting their local intensity distributions with two-dimensional Gaussian functions. Frame-wise displacement was calculated relative to the fitted position in the first frame. Rise and decay times were determined from the smoothed displacement-magnitude traces using the 10% to 90% and 90% to 10% crossing intervals, respectively. Detailed procedures for decorrelation analysis, Gaussian-based feature tracking, and kinetic quantification are provided in **Supplementary Note 2**.

### Preparation, staining, and imaging of mouse primary cortical neurons

MatTek glass-bottom 35 mm dishes (MatTek Corporation, P35G-1.5-20-C) were prepared as follows: glass-bottom dishes were cleaned for 15 min in 1 M HCl applied to the glass, then rinsed twice in sterile HBSS and twice in sterile distilled water. Dishes were coated with 10 µg/ml poly-L-lysine (MiliporeSigma, P5899) in Borate Buffer (0.01M, pH 8.5) for 10-16 h in 37 °C, followed by five 1 h washes with sterile water, then left overnight incubating in MEM (Corning, 10-010-CV) supplemented with 10% FBS (Life Technologies, A3160402). Cortices isolated from C57BL6 E17 mouse embryos were dissected and dissociated with trypsin at 37 °C (Thermo Fisher Scientific, 15050-065), according to the standard protocol from the Bassell lab^76^. Cells were plated at a density of 160.000 per dish. After 2 h, medium was replaced with astrocyte-conditioned Neurobasal Plus Medium (Gibco, A3582901) with B-27 Plus (Gibco, A3582801) and GlutaMAX (Gibco, 35050061), prepared as described earlier^76^. Mouse care and embryonic neuron isolation were performed in adherence to policies and regulations from the Institutional Animal Care and Use Committee (IACUC) at Emory University.

At DIV 4-5, standard culture medium was replaced with CO₂-independent Hibernate-E Low Fluorescence medium (BrainBits, HELF) with B-27 Plus and GlutaMAX. Experiments were performed on neurons at 8 and 16–21 days in vitro (DIV 8 and DIV 16–21). For dye loading, cells were incubated with 10 μM Fluo-4 AM (Thermo Fisher, F14201) in the maintenance medium for 45 min at 37°C. After three washes to remove excess dye, cells were left in HibE/B27+/GlutaMax, pre-equilibrated in a cell culture incubator for 1 to 2 hours. Cells were then incubated for an additional 30 min at 37 °C prior to data acquisition.

### Neuron Imaging experimental setup and procedures

To depolarize neurons, a custom field excitation probe was constructed following the general design presented previously^6,77^. A biological electronic stimulation unit (S88 Biological Stimulator; Grass Instruments) was coupled to the probe. To synchronize the electrical stimulation pulse and video capture, an sCMOS camera was used as a slave device to the stimulator unit and connected via hardware trigger inputs.

Neurons were placed in a stage-top incubator at 37 °C, 5% CO_2_, and 95% relative humidity. The excitation probe was placed in the imaging dish, immersing the platinum electrode lines in recording solution 1 mm above the neuron layer. Electrical stimulation consisted of 30 V pulses with durations of 3, 30, or 300 ms, as specified for each experiment. Candidate healthy neurons were selected based on qualitative evaluation by eye using the microscope eyepiece. After identifying a candidate neuron, an excitation pulse was triggered using the biological stimulator unit, which also triggered high-speed video capture from the sCMOS camera. SPAD image sequences were acquired independently using camera control software during the stimulation experiment. The images were captured using a dual-camera imaging setup (**Supplementary Fig. S1**). The sCMOS recording window was set long enough to capture the full temporal range of the SPAD camera. To align the two image streams, field-averaged fluorescence intensity traces were extracted from the sCMOS and SPAD recordings and temporally registered by cross-correlation. The resulting temporal offset was applied to match the SPAD BP sequence to the corresponding sCMOS frames for downstream analysis.

### Neuron calcium wave propagation analysis

The reconstructed QM and corresponding sCMOS image sequences were spatially registered by cross-correlation. Binary masks were manually delineated from time-averaged intensity projections to restrict the analysis to selected neuronal structures. For each pixel within the mask, the fluorescence trace was smoothed using a smoothing spline and fitted over the rising phase with a rectified-linear-unit model. The fitted breakpoint, corresponding to the beginning of the fluorescence increase, was taken as the response onset time and spatially mapped across the neuronal structure. The same analysis was applied to the registered QM and sCMOS recordings. A detailed description of the processing and fitting procedure is provided in **Supplementary Note 3**.

### Chemical and biological materials

The sources of the chemicals and biological materials used in the experiments, including company names and catalog numbers, are listed in **Supplementary Table S3**. No further authentication procedures were conducted after receiving the stock cell lines directly from the suppliers.

### Statistics and reproducibility

The fluorescence staining protocol was repeated at least three times for each experiment. During data acquisition, samples were imaged at least 5 times per experiment. Images with optimal fluorescence brightness were selected for the figures.

## DATA AVAILABILITY

Data underlying the results presented in this paper may be obtained from the corresponding author upon request due to the large file size. Source data are provided with this paper.

## CODE AVAILABILITY

The code package for the quanta microscopy is available as Supplementary Software. The code has been written in MATLAB 2024a (MathWorks) and Python 3.11. The latest version of the software will be made available at https://github.com/ShuJiaLab/QM upon publication.

## ACKNOWLEDGEMENTS

We acknowledge the support of the National Science Foundation grants 2145235 (S.J.), 2503686 (S.J.), 2607639 (S.J.); National Institutes of Health grant R35GM124846 (S.J.); and the Asan Foundation Biomedical Science Scholarship (J.Y.).

## AUTHOR CONTRIBUTIONS STATEMENT

J.Y. and S.J. conceived and designed the project. J.Y., K.H., and Y.H. contributed to the construction of the optical system. J.Y. performed imaging experiments, trained networks, and processed data. K.Y. and Z.L. helped with sample preparation and data analysis. A.J.K. prepared the mouse primary cortical neuron samples. W.Z. prepared the hiPSC-induced cardiomyocyte samples. C.X. and G.J.B. provided biological insights. S.J. supervised the overall project. J.Y. and S.J. wrote the manuscript with input from all authors.

## COMPETING INTERESTS STATEMENT

The authors declare no competing interests.

## Notes

### Competing Interest Statement

The authors have declared no competing interest.

## REFERENCES

1 Ettinger, A. & Wittmann, T. Fluorescence live cell imaging. Methods Cell Biol 123, 77–94 (2014). 10.1016/B978-0-12-420138-5.00005-7

2 Frigault, M. M., Lacoste, J., Swift, J. L. & Brown, C. M. Live-cell microscopy - tips and tools. J Cell Sci 122, 753–767 (2009). 10.1242/jcs.033837

3 Ni, Q., Mehta, S. & Zhang, J. Live-cell imaging of cell signaling using genetically encoded fluorescent reporters. FEBS J 285, 203–219 (2018). 10.1111/febs.14134

4 Laissue, P. P., Alghamdi, R. A., Tomancak, P., Reynaud, E. G. & Shroff, H. Assessing phototoxicity in live fluorescence imaging. Nat Methods 14, 657–661 (2017). 10.1038/nmeth.4344

5 Icha, J., Weber, M., Waters, J. C. & Norden, C. Phototoxicity in live fluorescence microscopy, and how to avoid it. Bioessays 39 (2017). 10.1002/bies.201700003

6 Mandracchia, B. et al. High-speed optical imaging with sCMOS pixel reassignment. Nat Commun 15, 4598 (2024). 10.1038/s41467-024-48987-7

7 Zhang, J. et al. Pixel-wise programmability enables dynamic high-SNR cameras for high-speed microscopy. Nat Commun 15, 4480 (2024). 10.1038/s41467-024-48765-5

8 Li, X. et al. Real-time denoising enables high-sensitivity fluorescence time-lapse imaging beyond the shot-noise limit. Nat Biotechnol 41, 282–292 (2023). 10.1038/s41587-022-01450-8

9 Sheneman, L., Balogun, S., Johnson, J. L., Harrison, M. J. & Vasdekis, A. E. Near-zero photon bioimaging by fusing deep learning and ultralow-light microscopy. Proc Natl Acad Sci U S A 122, e2412261122 (2025). 10.1073/pnas.2412261122

10 Zhang, G. et al. Bio-friendly long-term subcellular dynamic recording by self-supervised image enhancement microscopy. Nat Methods 20, 1957–1970 (2023). 10.1038/s41592-023-02058-9

11 Priessner, M. et al. Content-aware frame interpolation (CAFI): deep learning-based temporal super-resolution for fast bioimaging. Nat Methods 21, 322–330 (2024). 10.1038/s41592-023-02138-w

12 Chan, S. H. What does a one-bit quanta image sensor offer? IEEE Transactions on Computational Imaging 8, 770–783 (2022).

13 Fossum, E. R., Ma, J., Masoodian, S., Anzagira, L. & Zizza, R. The Quanta Image Sensor: Every Photon Counts. Sensors (Basel) 16 (2016). 10.3390/s16081260

14 Dutton, N. A., Gyongy, I., Parmesan, L. & Henderson, R. K. Single Photon Counting Performance and Noise Analysis of CMOS SPAD-Based Image Sensors. Sensors (Basel) 16 (2016). 10.3390/s16071122

15 Ma, J., Chan, S. & Fossum, E. R. Review of Quanta Image Sensors for Ultralow-Light Imaging. IEEE Transactions on Electron Devices 69, 2824–2839 (2022). 10.1109/ted.2022.3166716

16 Niclass, C., Rochas, A., Besse, P. A. & Charbon, E. Design and characterization of a CMOS 3-D image sensor based on single photon avalanche diodes. Ieee J Solid-St Circ 40, 1847–1854 (2005). 10.1109/jssc.2005.848173

17 Rochas, A. Single photon avalanche diodes in CMOS technology, EPFL, (2003).

18 Ulku, A. C. et al. A 512×512 SPAD Image Sensor with Integrated Gating for Widefield FLIM. IEEE J Sel Top Quantum Electron 25 (2019). 10.1109/JSTQE.2018.2867439

19 Ma, S. et al. Quanta burst photography. ACM Transactions on Graphics 39 (2020). 10.1145/3386569.3392470

20 Liu, Y. H., Kru, A., Basevi, H., Leonardis, A. & Jenkins, M.: 1-bit quanta video reconstruction by self-supervised photon location prediction. Adv Neur In (2024).

21 Yang, F., Lu, Y., Sbaiz, L. & Vetterli, M. in IS&T/SPIE Electronic Imaging. (SPIE).

22 Yang, F., Sbaiz, L., Charbon, E., Süsstrunk, S. & Vetterli, M. in 2009 IEEE 12th International Conference on Computer Vision Workshops, ICCV Workshops. 2212–2219.

23 Yang, F., Lu, Y. M., Sbaiz, L. & Vetterli, M. Bits from photons: oversampled image acquisition using binary Poisson statistics. IEEE Trans Image Process 21, 1421–1436 (2012). 10.1109/TIP.2011.2179306

24 Gnanasambandam, A. & Chan, S. H. HDR Imaging With Quanta Image Sensors: Theoretical Limits and Optimal Reconstruction. IEEE Transactions on Computational Imaging 6, 1571–1585 (2020). 10.1109/tci.2020.3041093

25 Chan, S. H. & Lu, Y. M. in 2014 IEEE Global Conference on Signal and Information Processing (GlobalSIP). 312–316.

26 Chan, S. H., Wang, X. & Elgendy, O. A. Plug-and-Play ADMM for Image Restoration: Fixed-Point Convergence and Applications. IEEE Transactions on Computational Imaging 3, 84–98 (2017). 10.1109/tci.2016.2629286

27 Chan, S. H., Elgendy, O. A. & Wang, X. Images from Bits: Non-Iterative Image Reconstruction for Quanta Image Sensors. Sensors (Basel) 16, 1961 (2016). 10.3390/s16111961

28 Chi, Y., Gnanasambandam, A., Koltun, V. & Chan, S. H. 122–138 (Springer International Publishing).

29 Choi, J. H., Elgendy, O. A. & Chan, S. H. in 2018 IEEE International Conference on Acoustics, Speech and Signal Processing (ICASSP). 6543–6547.

30 Seets, T., Ingle, A., Laurenzis, M. & Velten, A. in Proceedings of the IEEE/CVF Winter Conference on Applications of Computer Vision. 1945–1954.

31 Zhang, D., Lian, Q., Su, Y. & Ren, T. Dual-Prior Integrated Image Reconstruction for Quanta Image Sensors Using Multi-Agent Consensus Equilibrium. IEEE/CAA Journal of Automatica Sinica 10, 1407–1420 (2023). 10.1109/jas.2023.123390

32 Chennuri, P. et al. in Computer Vision – ECCV 2024: 18th European Conference, Milan, Italy, September 29–October 4, 2024, Proceedings, Part XL 152–171 (Springer-Verlag, Milan, Italy, 2024).

33 Garg, A., Ma, S. & Gupta, M. in Proceedings of the IEEE/CVF Conference on Computer Vision and Pattern Recognition. 19759–19770.

34 Seguret, M., Vermersch, E., Jouve, C. & Hulot, J. S. Cardiac Organoids to Model and Heal Heart Failure and Cardiomyopathies. Biomedicines 9 (2021). 10.3390/biomedicines9050563

35 Zhang, J. Z. et al. Protocol to measure contraction, calcium, and action potential in human-induced pluripotent stem cell-derived cardiomyocytes. STAR Protoc 2, 100859 (2021). 10.1016/j.xpro.2021.100859

36 Varzideh, F., Mone, P. & Santulli, G. Bioengineering Strategies to Create 3D Cardiac Constructs from Human Induced Pluripotent Stem Cells. Bioengineering (Basel) 9, 168 (2022). 10.3390/bioengineering9040168

37 Louch, W. E. et al. Reduced synchrony of Ca2+ release with loss of T-tubules-a comparison to Ca2+ release in human failing cardiomyocytes. Cardiovasc Res 62, 63–73 (2004). 10.1016/j.cardiores.2003.12.031

38 Kumar, S. et al. High-speed 2D and 3D fluorescence microscopy of cardiac myocytes. Opt Express 19, 13839–13847 (2011). 10.1364/OE.19.013839

39 Hinsdale, T. A., Stallinga, S. & Rieger, B. High-speed multicolor structured illumination microscopy using a hexagonal single mode fiber array. Biomed. Opt. Express 12, 1181–1194 (2021). 10.1364/BOE.416546

40 Bazmi, M. & Escobar, A. L. Autonomic Regulation of the Goldfish Intact Heart. Front Physiol 13, 793305 (2022). 10.3389/fphys.2022.793305

41 Denis, A. et al. Diagnostic value of isoproterenol testing in arrhythmogenic right ventricular cardiomyopathy. Circ Arrhythm Electrophysiol 7, 590–597 (2014). 10.1161/CIRCEP.113.001224

42 Zou, Y. et al. Isoproterenol activates extracellular signal-regulated protein kinases in cardiomyocytes through calcineurin. Circulation 104, 102–108 (2001). 10.1161/hc2601.090987

43 Ling, Z. et al. Multiscale and recursive unmixing of spatiotemporal rhythms for live-cell and intravital cardiac microscopy. Nat Cardiovasc Res 4, 637–648 (2025). 10.1038/s44161-025-00649-7

44 Grienberger, C. & Konnerth, A. Imaging calcium in neurons. Neuron 73, 862–885 (2012). 10.1016/j.neuron.2012.02.011

45 Yang, W. & Yuste, R. In vivo imaging of neural activity. Nat Methods 14, 349–359 (2017). 10.1038/nmeth.4230

46 Roome, C. J. & Kuhn, B. Simultaneous dendritic voltage and calcium imaging and somatic recording from Purkinje neurons in awake mice. Nat Commun 9, 3388 (2018). 10.1038/s41467-018-05900-3

47 Stringer, C., Michaelos, M., Tsyboulski, D., Lindo, S. E. & Pachitariu, M. High-precision coding in visual cortex. Cell 184, 2767–2778 e2715 (2021). 10.1016/j.cell.2021.03.042

48 London, M. & Hausser, M. Dendritic computation. Annu Rev Neurosci 28, 503–532 (2005). 10.1146/annurev.neuro.28.061604.135703

49 Higley, M. J. & Sabatini, B. L. Calcium signaling in dendrites and spines: practical and functional considerations. Neuron 59, 902–913 (2008). 10.1016/j.neuron.2008.08.020

50 Stuart, G. J. & Spruston, N. Dendritic integration: 60 years of progress. Nat Neurosci 18, 1713–1721 (2015). 10.1038/nn.4157

51 Bagheri, N., Carpenter, A. E., Lundberg, E., Plant, A. L. & Horwitz, R. The new era of quantitative cell imaging-challenges and opportunities. Mol Cell 82, 241–247 (2022). 10.1016/j.molcel.2021.12.024

52 Lemon, W. C. & McDole, K. Live-cell imaging in the era of too many microscopes. Curr Opin Cell Biol 66, 34–42 (2020). 10.1016/j.ceb.2020.04.008

53 Skylaki, S., Hilsenbeck, O. & Schroeder, T. Challenges in long-term imaging and quantification of single-cell dynamics. Nat Biotechnol 34, 1137–1144 (2016). 10.1038/nbt.3713

54 Hsieh, H. C. et al. Imaging 3D cell cultures with optical microscopy. Nat Methods 22, 1167–1190 (2025). 10.1038/s41592-025-02647-w

55 Mertz, J. Strategies for volumetric imaging with a fluorescence microscope. Optica 6, 1261–1268 (2019).

56 Hong, D. et al. Hyperspectral imaging. Nature Reviews Methods Primers 6, 19 (2026).

57 Cutrale, F. et al. Hyperspectral phasor analysis enables multiplexed 5D in vivo imaging. Nat Methods 14, 149–152 (2017). 10.1038/nmeth.4134

58 Barentine, A. E. S. et al. An integrated platform for high-throughput nanoscopy. Nat Biotechnol 41, 1549–1556 (2023). 10.1038/s41587-023-01702-1

59 Rees, P., Summers, H. D., Filby, A., Carpenter, A. E. & Doan, M. Imaging flow cytometry: a primer. Nat Rev Methods Primers 2, 86 (2022). 10.1038/s43586-022-00167-x

60 Lelek, M. et al. Single-molecule localization microscopy. Nat Rev Methods Primers 1, 39 (2021). 10.1038/s43586-021-00038-x

61 Ronceray, N. et al. Wide-field fluorescence lifetime imaging of single molecules with a gated single-photon camera. Light Sci Appl 14, 258 (2025). 10.1038/s41377-025-01901-2

62 Bucci, A. et al. 4D Single-particle tracking with asynchronous read-out single-photon avalanche diode array detector. Nat Commun 15, 6188 (2024). 10.1038/s41467-024-50512-9

63 Tseng, E. et al. Neural nano-optics for high-quality thin lens imaging. Nat Commun 12, 6493 (2021). 10.1038/s41467-021-26443-0

64 Barulin, A. et al. Dual-wavelength metalens enables Epi-fluorescence detection from single molecules. Nat Commun 15, 26 (2024). 10.1038/s41467-023-44407-4

65 Hoang, T. et al. 300-unit-per-second roll-to-roll manufacturing of visible metalenses. Nature 652, 1188–1194 (2026). 10.1038/s41586-026-10369-y

66 Morimoto, K. et al. Megapixel time-gated SPAD image sensor for 2D and 3D imaging applications. Optica 7, 346–354 (2020).

67 Na, N. et al. Room temperature operation of germanium-silicon single-photon avalanche diode. Nature 627, 295–300 (2024). 10.1038/s41586-024-07076-x

68 Shroff, H., Testa, I., Jug, F. & Manley, S. Live-cell imaging powered by computation. Nat Rev Mol Cell Biol 25, 443–463 (2024). 10.1038/s41580-024-00702-6

69 Wang, Z., Peng, Y., Fang, L. & Gao, L. Computational optical imaging: on the convergence of physical and digital layers. Optica 12, 113–130 (2025).

70 Defienne, H. et al. Advances in quantum imaging. Nature Photonics 18, 1024–1036 (2024).

71 Pearce, E., Nothlawala, F., Forbes, A. & Padgett, M. J. Quantum imaging with correlated photon pairs. Nature Reviews Methods Primers 6, 17 (2026).

72 Alieva, M., Wezenaar, A. K. L., Wehrens, E. J. & Rios, A. C. Bridging live-cell imaging and next-generation cancer treatment. Nat Rev Cancer 23, 731–745 (2023). 10.1038/s41568-023-00610-5

73 Zhu, Y., Moerner, W. E. & Qi, L. S. CRISPR-Cas-based live cell imaging of genome dynamics. Nat Rev Genet 27, 622–641 (2026). 10.1038/s41576-026-00949-z

74 Zuo, F. et al. Live-cell imaging of RNA dynamics using bright and stable fluorescent RNAs. Nat Protoc 21, 3765–3794 (2026). 10.1038/s41596-026-01343-z

75 Yoshimura, H. & Ozawa, T. Possible Direction of Drug Discovery Based on Single-Molecule Live Imaging. Annu Rev Pharmacol Toxicol 66, 227–240 (2026). 10.1146/annurev-pharmtox-062624-025717

76 Shi, L., Bagchi, P., Nguyen, G. N., Faundez, V. & Bassell, G. J. Protocol for identifying surface membrane proteins and their associated proteome from mouse cortical neuron cultures by in situ biotinylation. STAR Protoc 7, 104418 (2026). 10.1016/j.xpro.2026.104418

77 Stern, S., Rotem, A., Burnishev, Y., Weinreb, E. & Moses, E. External Excitation of Neurons Using Electric and Magnetic Fields in One- and Two-dimensional Cultures. J Vis Exp, e54357 (2017). 10.3791/54357

